# Discovering the Interactions between Circular RNAs and RNA-binding Proteins from CLIP-seq Data using circScan

**DOI:** 10.1101/115980

**Authors:** Bin Li, Xiao-Qin Zhang, Shu-Rong Liu, Shun Liu, Wen-Ju Sun, Qiao Lin, Yu-Xia Luo, Ke-Ren Zhou, Chen-Min Zhang, Ye-Ya Tan, Jian-Hua Yang, Liang-Hu Qu

**Author notes:** **Corresponding author:** Mailing address: Biotechnology Research Center, Sun Yat-sen University, Guangzhou, 510275, P. R. China, Jian-Hua Yang, Liang-Hu Qu. These authors contributed equally to this work (Bin Li, Xiao-Qin Zhang).

## Abstract

Although tens of thousands of circular RNAs (circRNAs) have been identified in mammalian genomes, only few of them have been characterized with biological functions. Here, we report a new approach, circScan, to identify regulatory interactions between circRNAs and RNA-binding proteins (RBPs) by discovering back-splicing reads from Cross-Linking and Immunoprecipitation followed by high-throughput sequencing (CLIP-seq) data. By using our method, we have systematically scanned ~1500 CLIP-seq datasets, and identified ~12540 and ~1090 novel circRNA-RBP interactions in human and mouse genomes, respectively, which include all known interactions between circRNAs and Argonaute (AGO) proteins. More than twenty novel interactions were further experimentally confirmed by RNA Immunoprecipitation quantitative PCR (RIP-qPCR). Importantly, we uncovered that some natural circRNAs interacted with cap-independent translation factors eukaryotic initiation factor 3 (eIF3) and N6-Methyladenosine (m^6^A), indicating they can be translated into proteins. These findings demonstrate that circRNAs are regulated by various RBPs, suggesting they may play important roles in diverse biological processes.

## Introduction

Circular RNAs (circRNAs) represent a group of RNA molecules with ends covalently linked in a circle^1-3^. Recent studies using high-throughput RNA-seq data analysis have illuminated circRNAs as a new and huge class of non-coding RNAs (ncRNAs) ubiquitous in eukaryotes with distinct sizes and sources^4-10^. However, it is unclear whether the majority of circRNAs represent only the side products of pre-mRNA splicing or are produced in a regulated manner to affect specific cellular functions^1-3^.

The control and function of circRNAs are governed by the specificity of RNA-Binding Proteins (RBPs)^11-13^. One emerging theme is that circRNA can act as a sponge to titrate miRNA and guides the function of the Argonaute (AGO) effector protein^4,7^. For example, brain-related circRNA ciRS-7/CDR1-AS harbors dozens of conserved miR-7 seed matches and AGO binding sites^4,7^. Moreover, it has been proposed that circRNAs could sequester RBPs or as the scaffold of RBPs, and therefore could be involved in intracellular transport of RBPs or RNAs^14^. On the other hand, although recent reports have indicated that short intronic repeat sequences facilitate circular RNA production^5,6,8,15^, splicing-related RBPs may participate in regulating circRNA biogenesis^15,16^. Indeed, production of many circRNAs was recently shown to be regulated by the Muscleblind and Quaking (QKI) splicing factors^16^. However, for the majority of circRNAs, the mechanism underlying their interaction with RBPs remains unknown.

Recent advances in high-throughput sequencing of RNAs isolated by crosslinking and immunoprecipitation (CLIP-seq) have provided powerful ways to identify RBP-associated RNAs and map these interactions in the genome^11–13,17–19^. The application of CLIP-seq methods in AGO and other RBPs have revealed that various types of linear RNA genes, such as mRNAs, lncRNAs and pseudogenes, can be regulated by miRNAs and RBPs.^10–12,20^. While an increasing number of RBPs have been explored using CLIP technologies, sequencing reads unaligned to genomes and known linear transcripts have been routinely discarded and excluded from further analysis. Moreover, CLIP-seq sequences were typically short (18-50 nt), existing circRNA tools required sequencing reads with long length (>100 nt) and are therefore limited to the detection of circRNAs from long RNA-seq datasets^1,3,5,7^.Therefore, it remains unknown whether the unmapped sequencing reads generated by various CLIP-seq technologies can be used to identify circRNAs as well as their interactions with RBPs.

To repurpose the publically available CLIP-seq datasets to interrogate the interactions between circRNAs and RBPs, we developed a computational algorithm, circScan, to reliably identify back-splicing junctions of circRNAs. We identified thousands of circRNAs interacting with RBPs from CLIP-seq datasets in human and mouse genomes. Using RNA Immunoprecipitation quantitative PCR (RIP-qPCR), we validated a subset of circRNA candidates interacted with cap-independent translation factors eukaryotic initiation factor 3 (eIF3) and N6-Methyladenosine (m^6^A)^21,22^, as well as microRNA effector protein AGO. Furthermore, polysome experiments and transcriptome-wide ribosome profiling analysis indicate, to the best of our knowledge for the first time, the translational potentials of circRNAs. Collectively, our approaches reveal extensive and complex circRNA-RBP interaction maps in human and mouse, and provide robust tool for further elucidating the functions as well as mechanisms of circRNAs.

## Results

### CircScan to precisely identify junction reads from CLIP-seq datasets

To search circRNAs in a more sensitive way, we designed a novel algorithm, circScan, which is specific for the screening of circRNAs from short CLIP-seq reads. In contrast to the known circRNA tools that mapped unaligned reads to genome, circScan searched for donor and acceptor anchors in unaligned CLIP-seq reads and stitched to produce full read alignments with back-splicing signals. The whole procedure is outlined in Figure 1 and described in detail in Materials and Methods. To detect whether the two known functional circRNAs, including brain-related CDR1-A S, and ubiquitously expressed circHIPK3, existed in short CLIP-seq data, we applied our method to a published AGO CLIP-seq dataset in human brain tissues. This analysis identified 471 circRNAs interacting with AGO protein (**Supplementary Tables 1**), including known CDR1-AS^4,7^ (**Supplementary Fig. 1a**) and circHIPK3^23^ circRNAs (**Supplementary Fig. 1b**). We found that 406 of 471 (86%) circRNAs were also overlapping with circRNAs identified from long RNA-seq data in human brain tissues (**Supplementary Fig. 1c**), and all host genes of candidate circRNAs were significantly enriched at neuronal gene ontology (GO) term (**Supplementary Fig. 1d**), suggesting that high accuracy of our predictions for short CLIP-seq reads.

**Figure 1.**
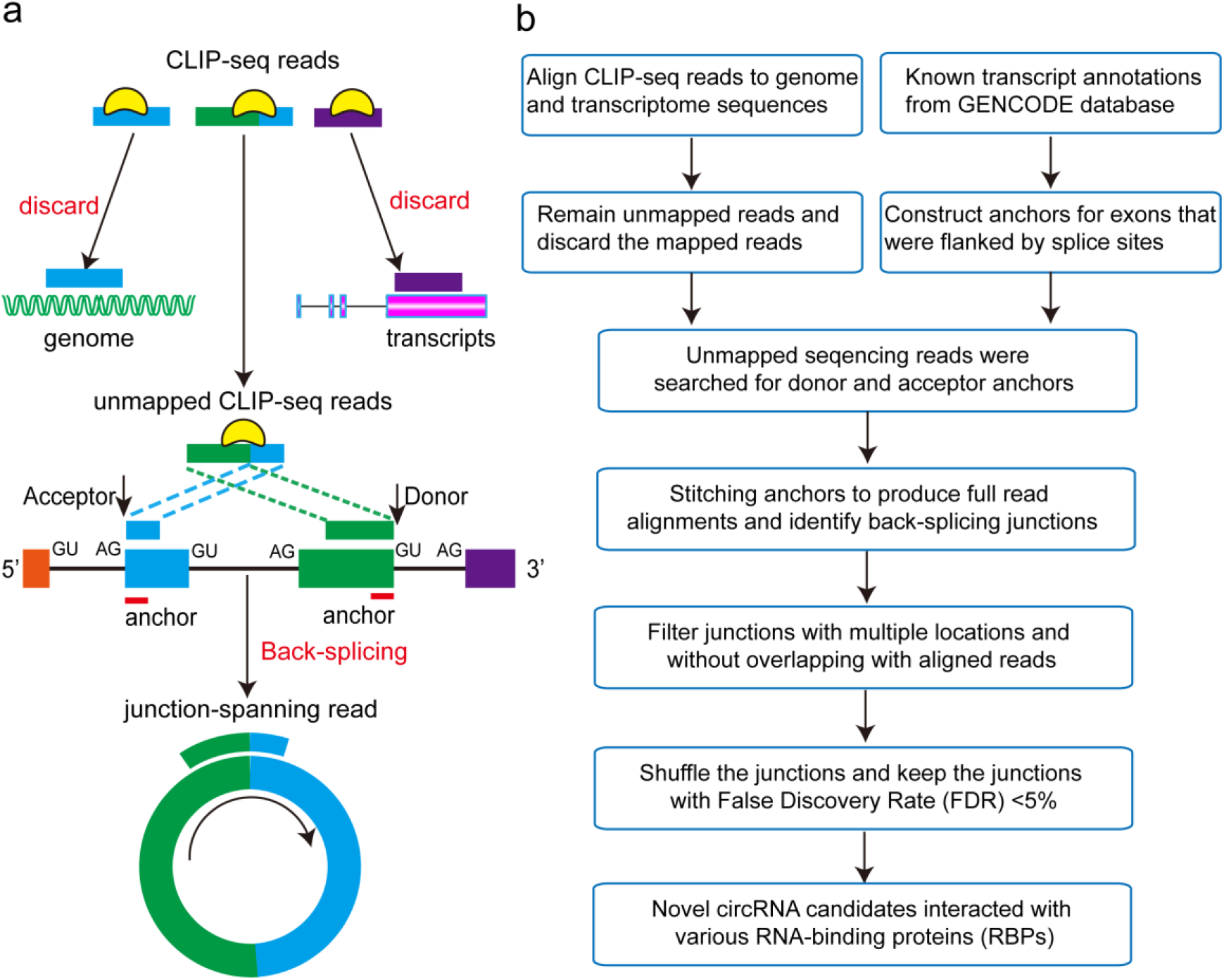
circScan algorithm workflow. **a**, Two-part alignments identified junction-spanning CLIP-seq reads indicative of circRNAs. Exons are colored, and donor (GU) and acceptor (AG) signals and anchors at splice sites are indicated. **b**, The computational workflow developed to identify and quantify circRNAs from short-read CLIP-seq data.

### Experimental validation of interactions between circRNAs and Ago proteins

To verify that circScan identified bona fide circRNAs rather than false positives, we experimentally tested our circRNA predictions in HEK293 cells. Head-to-tail splicing was assayed by qPCR after reverse transcription, with divergent primers. Predicted head-to-tail junctions of 6 out of 6 randomly chosen circRNAs (Fig. 2a) could be validated. To rule out the possibility that these head-to-tail splicing are produced by trans-splicing or genomic rearrangements as well as potential PCR artefacts, we quantified the RNase R resistance of these 6 candidates with confirmed head-to-tail splicing by qPCR. All of them were resistant to the RNase R (Fig. 2b). Furthermore, to determine whether these circRNAs can serves as a binding platform for AGO proteins, we conducted AGO2 immunoprecipitation in HEK293T cells (**Supplementary Fig. 2a**), and CDR1-AS and known miRNAs were selected as positive controls (**Supplementary Fig. 2b**). By RIP-qPCR assay, six endogenous circRNAs pulled down by AGO2 were specifically enriched more than 10-fold as compared to those pulled down by normal IgG protein (Fig. 2c). Taken together, these results further indicate the advantage of circScan software, and that many novel circRNAs may act as miRNA sponges.

**Figure 2.**
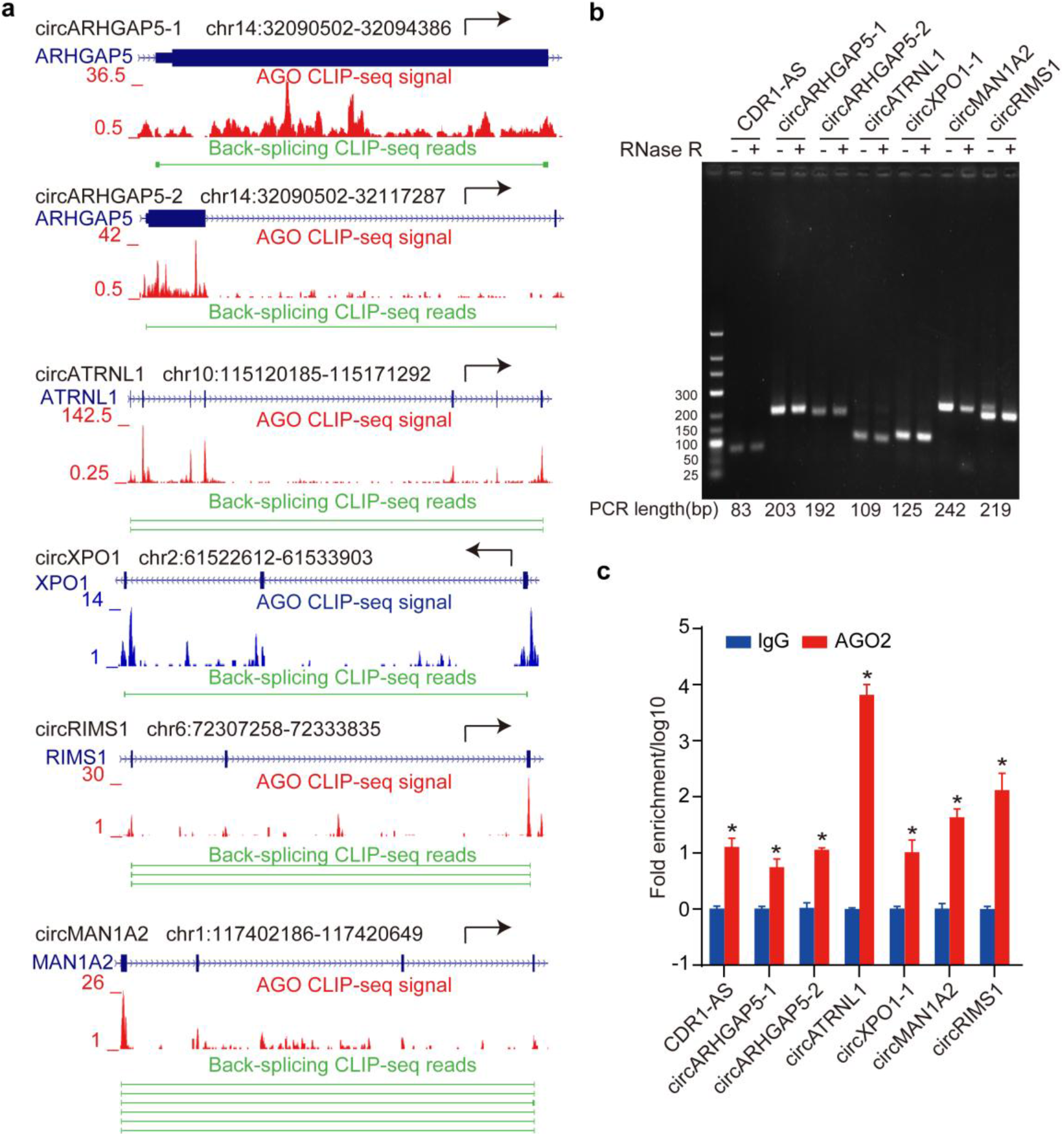
Experimental validation of the interactions between circRNAs and AGO protein. **a**, Selected randomly human circRNAs interacting with **AGO** protein for experimental validation. **b**, Confirmation and characterization of selected AGO2 associated circRNAs in HEK239T cells using RNase R followed by RT-PCR with divergent primers. And the PCR product size was labeled at the bottom of the figure. CDR1-AS was used as positive control. **c**, Validation of the selected circRNAs interacting with AGO2 by RIP-qPCR. RIP was performed from HEK293T cells using anti-AGO2 and IgG. Data are presented with respect to IgG that is set to a value of 1. The relative immunoprecipitate (IP)/IgG ratios (log10) are plotted. Data are shown as mean±SEM from three independent experiments. * p<0.05(student’s t-test).

### Transcriptome-wide search of circRNAs with circScan identifies thousands of circRNA-RBP interactions in the human and mouse genome

To investigate the interactions between various RBPs and circRNAs, we applied the program to the human genome for junction reads in published RBP CLIP-seq data sets across various CLIP methods, including HITS-CLIP, PAR-CLIP, eCLIP, iCLIP and CLEAR-CLIP etc. Our circScan algorithm confidently detected 12450 novel interactions between 106 RBPs and 8212 circRNAs (**Supplementary Table 2**). Numbers of circRNA-RBP interactions varied strongly across RBPs, presumably due to the differences in experiment number for each RBP (Fig. 3a, **Supplementary Table 3**). For example, we found that the top one IGF2BP2 RBP bound to 2139 novel circRNAs (**Supplementary Table 3**). While the majority of head-to-tail junctions were supported by few reads (one to three), we also detected 478 human circRNAs supported by more than 10 junction-spanning reads (**Supplementary Fig. 3a, Supplementary Table 2**). All circRNAs were usually composed of protein-coding exons, with a clear preference for coding sequence (CDS) exons (Fig. 3b). Although the majority of circRNAs were bound by only one or two RBPs, 58 circRNAs, including CDR1as ^4,7^, were found to harbore binding sites of over 10 RBPs (Fig. 3c, **Supplementary Table 4**). This finding suggested the diverse roles of circRNAs in biological processes when accompanied with different RBPs.

**Figure 3.**
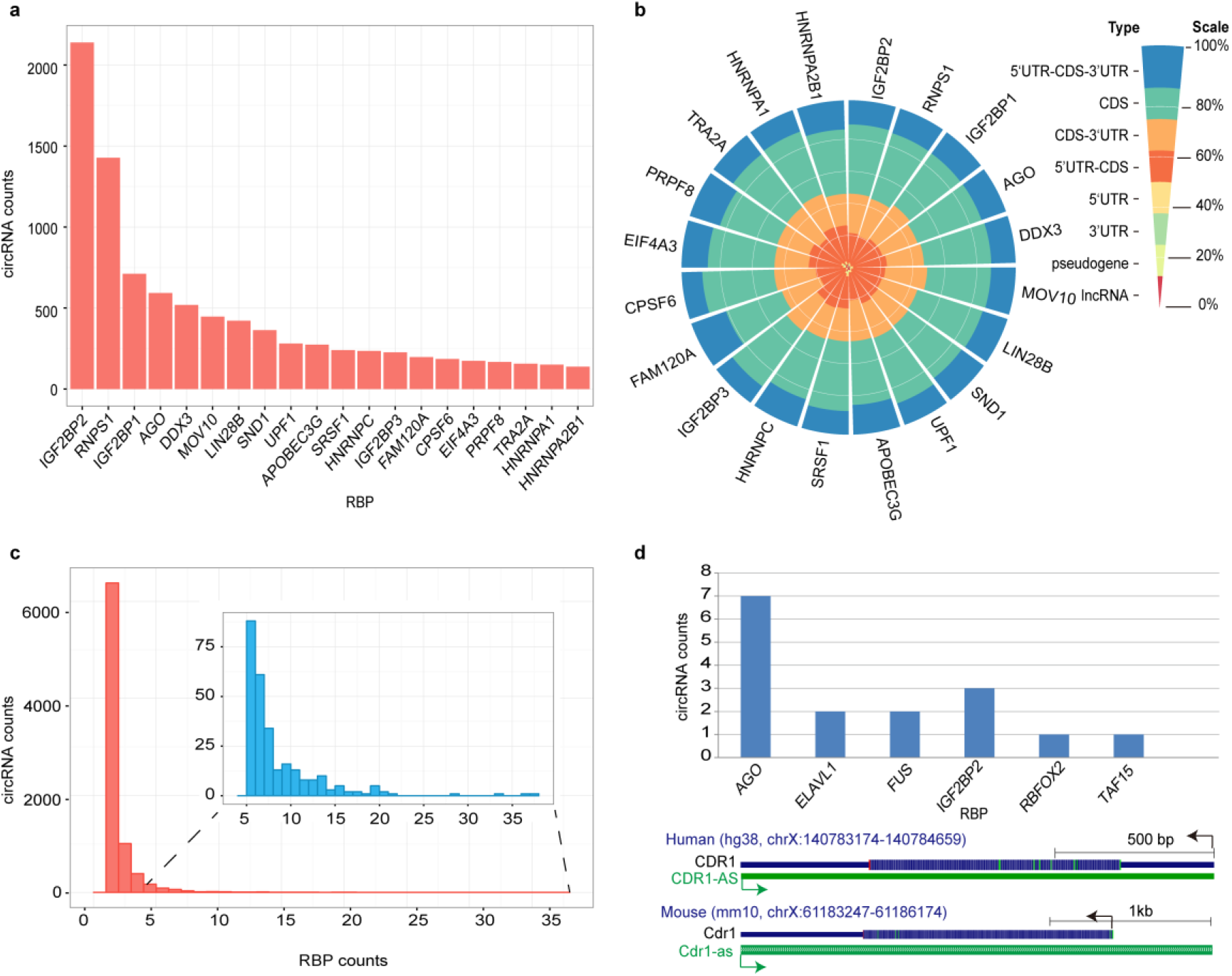
Interactions between RBPs and circRNAs identified by circScan. **a**, The distribution of circRNAs bound by top 20 RBPs. **b**, The genomic distributions of circRNAs bound by various RBPs. **c**, The distribution of circRNAs bound by different numbers of RBPs. **d**, the conserved interactions between RBPs and circRNAs and an example derives from human CDR1-AS and mouse Cdr1-as loci.

To determine the evolutionary conservation and consequent functional importance of circRNAs, the human and mouse circRNA interactomes were compared. We applied circScan to CLIP-seq datasets derived from mouse, and discovered 1094 novel interactions between 36 RBPs and 918 circRNAs (**Supplementary Table 5**). The distributions of the mouse circRNA-RBP interactions are similar with that in human genome (**Supplementary Fig. 3b-d**). We next systematically assessed the extent of circRNAs conservation by using whole-genome alignments to identify the regions of the mouse genome that corresponded to the human circRNAs. Of the 8,212 circRNAs interacting with RBPs in human, 2091 could be reliably mapped to known and orthologous mouse circRNAs downloaded from public databases^24^ (**Supplementary Fig. 3e**). However, due to the differences in the number of CLIP-seq experiments and RBPs between human and mouse, we only identified 16 evolutional conservation of circRNA-RBP interactions between 6 RBPs and 16 circRNAs (Fig. 3d). For example, the interactions between CDR1-AS and AGO or FUS proteins are conserved in the human and mouse genome (Fig. 3d).

### Exploring translational potentials of endogenous circRNAs interacting with eIF3 and m^6^A

Given that we observed that majority of circRNAs were derived from CDS exons, we surveyed their potential to be translated into proteins. While circRNAs possess neither a 5’ nor a 3’ end, we postulated that circRNAs with translational potentials may interact with cap-independent translation initiation-associated factors^21,22^, such as eIF3 and m^6^A. Using circScan to eIF3 CLIP-seq data, we identified 20 circRNAs interacted with eIF3 (**Supplementary Table 6**) and confirmed all of them were resistant to the RNase R (Fig. 4a). With the exception of four circRNAs (circPCMT1, circEARS2, circCD276 and circABCA3) with multiple PCR bands, sixteen endogenous circRNAs were selected to eIF3 RIP-qPCR assays (**Supplementary Fig. 4a**) and three genes (MYC, GAPDH, MALAT1) were selected positive controls (**Supplementary Fig. 4b**). We found that they were all specifically enriched more than 10-fold as compared to those pulled down by normal IgG protein (Fig. 4b, **Supplementary Table 6**).

**Figure 4.**
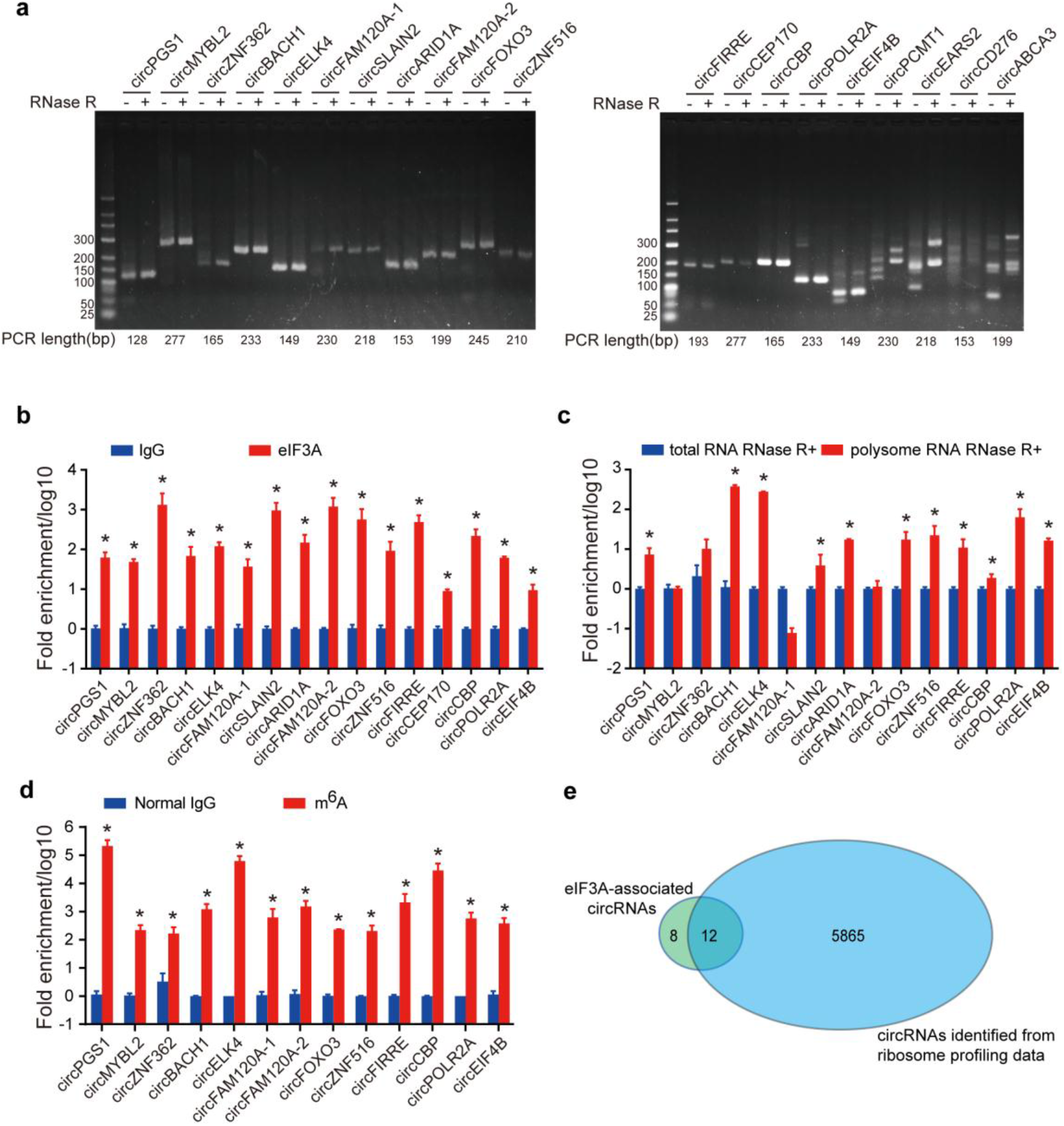
Translational potentials of natural circRNAs. **a**, Confirmation and characterization of selected eIF3A associated circRNAs in HEK239T cells using RNase R followed by RT-PCR with divergent primers. **b**, Validation of the selected circRNAs interacting with eIF3A by RIP-qPCR. RIP was performed from HEK293T cells using anti-eIF3A and IgG. Data are presented with respect to IgG that is set to a value of 1. Data are shown as mean±SEM from three independent experiments. * p<0.05(student’s t-test). **c**, qPCR detection of eIF3A associated circRNAs in RNase R treated Polysome RNA and total RNA. Data are presented with respect to RNase R treated total RNA that is set to a value of 1. Data are shown as mean ± SEM from two independent experiments. * p<0.05(student’s t-test). **d**, MeRIP-qPCR analysis of RNase R treated RNA isolated from HEK239T cells and immunoprecipitated with anti-m6A or IgG. Data are presented with respect to IgG that is set to a value of 1. Data are shown as mean±SEM from three independent experiments. * p<0.05(student’s t-test). **e**, The overlap between eIF3A-associated circRNAs and circRNAs identified from ribosome profiling data.

We next explored whether these circRNAs might be associated with m^6^A. We also found 12 circRNAs interacting with eIF3 also can be bound by m^6^A reader YTHDFs by analyzing public RIP-seq (non-polyA) data (**Supplementary Table 6**). We further examined all circRNAs using an antibody against m^6^A (**Supplementary Fig. 4c**) in the HEK293T cells and detected thirteen circRNAs bound to m^6^A (Fig. 4c, **Supplementary Table 6**). The discovery of m^6^A in circRNAs also suggests that this class of ncRNAs is regulated by diverse RNA modifications.

After the discovery that many circRNAs are associated with eIF3 and m^6^A, we next detected whether these circRNAs interacted with translation-related ribosomes. We isolated polysome-bound fractions by ultracentrifugation and assayed the relative quantity of circRNAs by qPCR. Eleven circRNAs were significantly enriched in the ribosome-bound fraction (Fig. 4d, **Supplementary Table 6**), indicating that these circRNAs may be translated.

To further investigated the association of circRNAs with ribosomes, we applied our circScan to ribosome-protected fragments (RPFs) generated by ribosome profiling and detected 5877 circRNAs (Fig.4e, **Supplementary Table 7**). Of the above-mentioned 20 circRNAs interacting with eIF3, 12 could be found in the ribosome profiling datasets, providing another evidence for circRNA translation.

## Discussion

Lack of knowledge regarding the interacted RBPs of circRNAs was one of the major factors that limited understanding of their roles in various biological processes. CircScan provides a valuable tool to bridge this gap and allows global maps of the RBP-circRNA interactions to resolve some obstacles that have arisen in the efforts to understand circRNA function. In this study, majority of the random-selected circRNAs which can interact with three different antibodies (AGO, eIF3 and m^6^A) can be confirmed by RIP-qPCR, demonstrating that our circScan software may help users to gain RBP-circRNA interactions of high-confidence. Importantly, we firstidentify circRNAs as a new class of m^6^A-containing RNAs, and provid initial evidences for that circRNAs might be translated into proteins.

Although more than 100,000 circRNAs have been identified in diverse tissues or cell lines, only a few of them have been confirmed with biological functions^4–9^. This disequilibrium has raised the debate in the field concerning the function of circRNA.

For example, one study suggested that the majority of circRNAs are mere by-products of pre-mRNA ^25^, and many studies have suggested that circRNAs may have biological functions based on the observations that the conservation of circRNA sequences across vertebrates and the developmental regulation of circRNAs^7–9,26,27^. The direct evidence that determined the biological functions of one circRNA is what RBP interacted with it^11, 12^. In this study, more than 10,000 interactions identified from >200 different RBPs in human and mouse strongly support a large number of circRNAs have biological functions. Moreover, by combining binding maps for all RBPs, we found that one circRNA was often bound by multiple RBPs (Fig. 3c). This is consistent with the compelling idea that circRNAs can serve as scaffolds that assemble multiple relevant RBPs to regulate gene expression. Notably, considering that we only detected the head-to-tail junction bound by RBPs, the study might miss the sequences outside the junction, and the number of functional circRNAs identified here may be underestimated.

While protein-encoding RNA circles are well-established as viroids ^28^, it is not clear whether endogenous circRNAs in eukaryotic genomes can be translated into proteins. The discovery that the synthesized circRNAs can be translated into proteins in human cells has increased the likelihood that some endogenous circRNAs can function as protein-coding RNA transcripts^29,30^. Here we provide initial evidence for the translational potentials of circRNAs by analyzing the eIF3 and m^6^A CLIP-seq data. Given the recently established fact that m^6^A promotes cap-independent translation in linear protein-coding genes and requirement of a novel m^6^A reader eIF3^21,31^, it seems reasonable to speculate that the translation of circRNAs also might be drived by m^6^A modifications. The potentials of translation is further confirmed in both ribosome profiling datasets and in polysome experiments (Fig. 4c, 4e). In addition, ~5800 circRNAs were discovered by applied our methods to ribosome profiling data and further reveal much more circRNAs might be translated into proteins.

In summary, we developed a novel algorithm for identifying interactions between circRNAs and RBPs through repurposing CLIP-seq datasets. Our study opens new avenues for leveraging publicly available genomic data to study the functions and mechanisms of circRNAs.

## Methods

**Cell culture**HEK293T and Hela cells were obtained from the Shanghai Institute of Cell biology, Chinese Academy of Science. HEK293T and Hela were cultured in DMEM (Gibco) supplemented with 10% fetal bovine serum (FBS, Gibco) and 1% penicillin–streptomycin solution (Gibco) at 37°C and 5% CO_2_. Cells were free of mycoplasma contamination based on MycoBlue Mycoplasma Detector (Vazyme) and STR profiling was used to authenticate the cell lines (Cellcook Biotech Co., Ltd, Guangzhou).

### RNA extraction, RT-PCR and Quantitative real-time PCR

Total RNAs from cultured cells were extracted with TRIzol reagent (Invitrogen) according to the manufacturer’s instruction. The RT-PCR for circRNAs and mRNAs were carried out with 5X All-In-One RT MasterMix (with AccuRT Genomic DNA Removal Kit) (abm). For miRNA reverse transcription, DNase I-digested RNA was reverse transcribed using a Mir-X miRNA First-Strand Synthesis Kit (Takara). Quantitative real-time PCR (qPCR) was performed using PowerUp SYBR Green Master Mix (A Δ BI) with three biological replicates according to the manufacturer’s suggestions. The comparative Ct Method (ΔCT Method) was used to determine the relative expression levels of genes. In particular, the divergent primers annealing at the distal ends of circRNA were used to determine the abundance of circRNA. Primers for qPCR used were listed in **Supplementary Table 8.**

### RNA-seq

RNA-sequencing was performed by Annoroad Gene Technology Co., Ltd, Beijing, China. Briefly, 3 μg of extracted RNA from HEK293T or Hela cells was used as input for the Illumina TruSeq^®^ Stranded Total RNA HT/LT Sample Prep Kit following depletion of ribosomal RNA using Ribo-Zero rRNA Removal Kit (Epicentre). Libraries were pooled and sequenced on an Illumina X-ten PE150 platform. A minimum of 300 M reads were generated per sample.

### Integration of public high-throughput sequencing data sets

High-throughput CLIP-seq, ribosome profiling and RIP-seq sequencing datasets were retrieved from the Gene Expression Omnibus (GEO), Sequence Read Archive (SRA)^32^ and ArrayExpress. The eCLIP datasets were downloaded from ENCODE websites. The datasets used in the study were listed in the **Supplementary Table 9.**

### Sequencing data processing

Barcodes or 3’-adapters of raw sequencing data were clipped using the FASTX-toolkit software (version 0.0.13). We set a read quality for each data set, retaining only those reads with a quality score above 20 in 80% of their nucleotides, and we also restricted the read length to 15 nucleotides (nt) after adapter trimming. Finally, we collapsed identical reads with same barcodes into one read for minimizing PCR duplicates.

### Algorithm description

The core circScan algorithm workflow was shown using a schematic diagram in Figure 1. The detailed information for each step was described as follow.

#### 1. Mapping CLIP-seq reads to genome and transcriptomes

All unique CLIP-seq reads without adapters and barcodes in each sample were mapped to the human and mouse genomes and transcriptomes using Bowtie 2.0 (parameters: -D 200 -R 3 -N 0 -L 15 -i S,1,0.5 --score-min=C,-16,0), and then converted alignment SAM format into the sorted BAM format. The aligned reads were discarded and the remaining unmapped CLIP-seq reads were used to identify circRNAs.

#### 2. Identifying circRNAs from high-throughput CLIP-seq datasets

For each transcript annotated in GENCODE for human (v25) and mouse (M10), we generated “anchors”, defined as 10 nt exon sequence that were flanked by GU/AG splice sites from each transcript. We called the anchor as “donor anchor” if it was near the donor site (GU) and “acceptor anchor” if it was near the acceptor site (AG). Unmapped sequencing reads were searched for donor and acceptor anchors, respectively. Reads containing both donor and acceptor anchors were extended to produce full read alignments and to determine whether they were back-splicing junctions. We chose the best alignment for each read using a scoring schema, which are described as follows: (i) match bonus 1, (ii) mismatch penalty 1 and (iii) sequencing reads of PAR-CLIP samples contain characteristic T to C mutations caused by crosslinking of photo-reactive nucleoside 4-thiouridine. Therefore, the T to C mutation was regarded as match in PAR-CLIP data and bonus 1. Junction reads with score (>=20) were further filtered with following steps. (i) With unique genomic location; (ii) to guarantee the circRNAs have expression, the donor and acceptor sites located within the head-to-tail junction must be overlapped with at least one aligned read; (iii) to filter out unreliable mappings, junction sequence was shuffled 1000 times and scored by the above-mentioned scoring schema, we kept the junctions with false discovery rate (FDR) <0.05 by defined as the number of shuffle reads with alignment score >= original score divide by total shuffle times.

#### 3. Annotation and filter of circRNAs

All gene annotations of human (GENCODE V25) and mouse genome (GENCODE M10) were downloaded from GENCODE website. All candidate circRNAs were annotated using above-mentioned annotation data sets, and only the circRNAs with the precise positions of downstream donor and upstream acceptor splice sites of known genes were kept.

### RNase R treatment

For RNase R treatment, RNAs were incubated with or without 5 U RNase R (Epicentre) for 3 h at 37°C and the resulting RNA was subsequently purified using TRIzol.

### Native RNA immunoprecipitation

RIP experiments were carried out in native conditions as described^33^. Briefly, approximately 1 × 10^7^ cells were pelleted and re-suspended with an equal pellet volume of ice-cold Polysomal Lysis Buffer (10 mM HEPES, pH 7.0, 100 mM KCl, 5 mM MgCl2, 0.5% NP40, 1mM DTT, 100 U/mL RNase Inhibitor (Takara), 1×Protease Inhibitor Cocktail (Roche) and 0.4 mM RVC (NEB) and lysate were passed through a 26 gauge needle 6 times and incubated on ice for 15 min. After the lysates were centrifuged at 15000 g for 15 min, the supernatants were pre-cleared with Dynabeads Protein G (Invitrogen). Then the lysate was diluted in NT2 buffer (50 mM Tris, pH 7.4, 150 mM NaCl, 1 mM MgCl_2_, 0.05% NP40, 1mM DTT, 100 U/mL RNase Inhibitor (Takara), 1 × Protease Inhibitor Cocktail (Roche) and 20 mM EDTA). One hundredth of the supernatant was saved as input and the rest lysates were used for RIP with 5 μg of various antibodies at 4°C overnight. On the next day, the RNA-Protein/antibody complex was precipitated by incubation with Dynabeads Protein G at 4°C for 3 h. The beads were washed five times with NT2 buffer. A quarter of the RIP material was used for Western blot and the rest was used for RNA extraction with TRIzol. The fold enrichment of RNAs was detected by qPCR assay. Antibodies directed against AGO2, eIF3A and normal mouse and rabbit IgG were respectively from Abnova H00027161-M01, Abcam ab86146, Millopore #12-371 and CST #2729.

### MeRIP-qPCR of circRNA

CircRNAs were isolated from HEK293T total RNA with DNase I digestion and RNase R treatment. 11 p μ g RNase R treated RNA were pre-cleared with Dynabeads Protein G (Invitrogen) in 1.1 mL 1 × m6A IP buffer(10 mM Tris-HCl (pH7.5), 150 mM NaCl, 0.1% NP-40) supplemented with 200U RNase Inhibitor (Takara), 1mM DTT and 2mM RVC(NEB) for 1 h at 4°C. After pre-clear, 50 pμ L supernatant was saved as input and the rest supernatant were used for MeRIP with 5 pμ g of anti-m6A (Abcam, ab151230) and Normal rabbit IgG (CST, 2729) at 4°C overnight, respectively. Then the RNA-antibody complex was precipitated by incubation with Dynabeads Protein G at 4°C for 3 h. Samples were washed four times with 1× m6A IP buffer, and RNA was eluted from the beads by 300 pμ L Elution Buffer (5 mM Tris-HCl (pH7.5), 1 mM EDTA pH 8.0, 0.05% SDS) supplemented with 8.4 pμ g Proteinase K (Beyotime) at 50°C for 90 min. Following phenol extraction and ethanol precipitation, the input and eluted RNA were reverse transcribed with random hexamers and enrichment was determined by qPCR.

### m6A dot-blotting

For m6A dot-blot analysis, HEK239T DNase I-digested total RNA, MeRIP Input, anti-IgG anti-m6A, DEPC-treated water and poly(A) RNA (2 pμ L) were spotted onto Hybond N+ nylon membranes (Amersham). After crosslinking at 120,000 pμ J/cm^2^ at 254 nm using HL-2000 HybriLinker (UVP, Inc.), the membrane was blocked with 5% non-fat milk in TBST(pH7.6, 20 mM Tris-HCl, 150 mM NaCl, 0.05 % Tween-20) for 1 h at room temperature and then incubated with m6A antibody (Abcam, ab151230) overnight at 4 °C. After washing with TBST, membranes were incubated with HRP-linked anti-rabbit IgG (CST, 7074) for 1 h at room temperature. Then the membrane was exposed to x-ray film by Immobilon Western Chemiluminescent HRP Substrate (Millipore, WBKLS0500).

### Polysome isolation

For Polysome isolation, cells were grown to ~90% confluence and four 150 mm dishes of HEK293T cells were used.Cycloheximide (MP Biomedicals) was added to a final concentration of 100 pμ g/mL and incubated for 10 min at 37°C. The cells were then rinsed twice with cold-PBS and lysed in 400 pμ L Mammalian Polysome Buffer (20 mM Tris, pH 7.5, 250 mM NaCl, 5 mM MgCl_2_, 1% Triton X-100, 1mM DTT, 10 U DNase I, 100 μg/mL Cycloheximide, 0.1% NP-40 and 500U/mL RNase Inhibitor). After passing the lysates through a 26 gauge needle 4 times, a 23 gauge needle 6 times and incubated on ice for 10 min,the lysates were centrifuged for 10 min at 20000 g at 4°C and a part of supernatant reserved for total RNA isolation. The rest supernatant was gently layered over the sucrose cushion (20 mM Tris, pH 7.5, 250 mM NaCl, 5 mM MgCl_2_, 1.461 M Sucrose, 1mM DTT, 100 pμ g/mL Cycloheximide and 500 U/mL RNase Inhibitor). After then, they were centrifuged at 4°C in the Beckman Optima L-100XP fixed-angle Type 100Ti rotor at 70000 rpm for 4 h, and the pellet containing polysomes was used for RNA extraction with TRIzol.

## Code Availability

circScan is implemented in C/C++ and Perl, and runs on Linux and Mac OS X. The circScan developed in this study are available on request to the corresponding authors.

## Author contributions

J.H.Y. conceived and designed the entire project. J.H.Y., and L.H.Q., designed and supervised the research. J.H.Y. wrote the computational software; B.L., X.Q.Z., S.L., W.J.S., K.R.Z. and J.H.Y. performed the genome-wide or transcriptome-wide data analyses; B.L., X.Q.Z., S.R.L., S.L., W.J.S., Q.L., Y.X.L., K.R.Z., C.M.Z., Y.Y.T. and J.H.Y. performed experiments and/or data analyses; B.L., X.Q.Z., J.H.Y. and L.H.Q. contributed reagents/analytic tools and/or rant support; B.L., X.Q.Z., S.R.L., S.L., W.J.S., Q.L., Y.X.L., K.R.Z., C.M.Z., Y.Y.T., J.H.Y. and L.H.Q. wrote the paper. All authors discussed the results and commented on the manuscript.

## Acknowledgments

This research is supported by the Ministry of Science and Technology of China, National Basic Research Program (No. 2011CB811300); the National Natural Science Foundation of China (No. 30900820, 31230042, 31370791, 31471223, 91440110); funds from Guangdong Province (No. S2012010010510, S2013010012457); The project of Science and Technology New Star in ZhuJiang Guangzhou city (No. 2012J2200025); Fundamental Research Funds for the Central Universities (No. 2011330003161070, 14lgjc18); China Postdoctoral Science Foundation (No. 200902348); This research is supported in part by the Guangdong Province Key Laboratory of Computational Science and the Guangdong Province Computational Science Innovative Research Team.

